# Rapid adaptation of SARS-CoV-2 in BALB/c mice: Novel mouse model for vaccine efficacy

**DOI:** 10.1101/2020.05.02.073411

**Authors:** Hongjing Gu, Qi Chen, Guan Yang, Lei He, Hang Fan, Yong-Qiang Deng, Yanxiao Wang, Yue Teng, Zhongpeng Zhao, Yujun Cui, Yuchang Li, Xiao-Feng Li, Jiangfan Li, Nana Zhang, Xiaolan Yang, Shaolong Chen, Guangyu Zhao, Xiliang Wang, Deyan Luo, Hui Wang, Xiao Yang, Yan Li, Gencheng Han, Yuxian He, Xiaojun Zhou, Shusheng Geng, Xiaoli Sheng, Shibo Jiang, Shihui Sun, Cheng-Feng Qin, Yusen Zhou

**Author notes:** Corresponding to: Shibo Jiang, Shihui Sun, Cheng-Feng Qin, Yusen Zhou. These authors contributed equally (Hongjing Gu, Qi Chen, Guan Yang, Lei He, Hang Fan, Yong-Qiang Deng).

## Abstract

Coronavirus disease 2019 (COVID-19) threatens global public health and economy. In order to develop safe and effective vaccines, suitable animal models must be established. Here we report the rapid adaption of SARS-CoV-2 in BALB/c mice, based on which a convenient, economical and effective animal model was developed. Specifically, we found that mouse-adapted SARS-CoV-2 at passage 6 (MACSp6) efficiently infected both aged and young wild-type BALB/c mice, resulting in moderate pneumonia as well as inflammatory responses. The elevated infectivity of MACSp6 in mice could be attributed to the substitution of a key residue (N501Y) in the receptor-binding domain (RBD). Using this novel animal model, we further evaluated the in vivo protective efficacy of an RBD-based SARS-CoV-2 subunit vaccine, which elicited highly potent neutralizing antibodies and conferred full protection against SARS-CoV-2 MACSp6 challenge. This novel mouse model is convenient and effective in evaluating the *in vivo* protective efficacy of SARS-CoV-2 vaccine.

**Summary:** This study describes a unique mouse model for SARS-CoV-2 infection and confirms protective efficacy of a SARS-CoV-2 RBD subunit vaccine.

## Introduction

A novel coronavirus (CoV), severe acute respiratory syndrome coronavirus 2 (SARS-CoV-2) that causes CoV disease 2019 (COVID-19), has infected millions of people with more than 139,000 deaths, leading to devastating damage to global public health and economy (*1–6*). Elderly COVID-19 patients generally have a higher case-fatality rate (CFR) than the younger patients with COVID-19, suggesting that aged people are more susceptible to SARS-CoV-2 infection (*7, 8*). SARS-CoV-2 is categorized in the same betacoronavirus genus as SARS-CoV that caused a global outbreak in 2003 (*9*). Yet, SARS-CoV-2 has superior capacity of human-to-human transmission (*7*). Still, no vaccines or therapeutic agents are currently available to combat the COVID-19 pandemic, thus calling for the development of safe and effective vaccines and therapeutics.

Among the four structural proteins of coronovirus, spike (S) plays the most important roles in viral infection, zoonotic and human-to-human transmission, and pathogenesis, as such it is a critical target for vaccine and therapeutic development (*10*). The S protein is composed of S1 and S2 subunits. Receptor-binding domain (RBD) of S1 subunit binds to angiotensin-converting enzyme 2 (ACE2) on the host cell, while S2 subunit mediates fusion between the viral and target cell membranes (*11*). Our previous studies have demonstrated that the RBDs of other CoVs, including SARS-CoV and Middle East respiratory syndrome coronavirus (MERS-CoV), contain major conformation-dependent neutralizing epitopes and that RBD-based subunit vaccines can elicit highly potent neutralizing antibodies in the immunized animals (*12–14*).

Small animal models, such as mice, provide reliable and convenient approaches to evaluate the in vivo efficacy of antiviral countermeasures, including vaccines and prophylactic/therapeutic agents. However, the wide-type mice are generally not susceptible to infection with human coronaviruses using human proteins as receptors. For example, MERS-CoV does not infect wildtype mice, but infects mice expressing human dipeptidyl peptidase 4 (DPP4) (*15*). It has been recently reported that human ACE2 (hACE2)-transgenic (hACE2-Tg) mice are susceptible to SARS-CoV-2 infection. However, the very limited availability of hACE2-Tg mice as well as the associated costs, including maintenance, has seriously hampered the development of COVID-19 vaccines and therapeutics (*16*).

Herein we report the identification of a mouse-adapted SARS-CoV-2 strain, MASCp6, after only 6 passages of SARS-CoV-2 in aged BALB/c mice. This mouse-adapted SARS-CoV-2 strain can infect and replicate efficiently in the lung and trachea of both young and aged BALB/c mice, causing damage and inflammatory responses resembling those in COVID-19 patients. Most important, our previously developed RBD-based SARS-CoV-2 subunit vaccine can elicit strong neutralizing antibody responses in vaccinated mice such that they are fully protected against SARS-CoV-2 (MASCp6) challenge. To the best of our knowledge, this is the first report of a small animal model for SARS-CoV-2 infection. It can be conveniently and economically used to evaluate the in vivo efficacy of vaccines, antibodies, and other antiviral agents against SARS-CoV-2 infection, robustly promoting the development of prophylactics and therapeutics for prevention and treatment of COVID-19.

## Results

### Generation of a mouse-adapted SARS-CoV-2 strain, MASCp6

To generate a mouse-adapted SARS-CoV-2 strain able to effectively replicate in wild-type mice, we serially passaged SARS-CoV-2 (BetaCoV/Wuhan/AMMS01/2020) in aged (9-month-old) BALB/c mice as previously described for SARS-CoV (*17*). We intranasally (i.n.) inoculated three aged mice with 7.2×10^5^ plaque forming unit (PFU) of SARS-CoV-2 and sacrificed them three days post-infection (dpi). We found that SARS-CoV-2 infected these mice and that the RNA copies (10^7.62^/mouse) in the lung homogenate were detected. We defined this homogenate as passage 0 (P0), and used it to infect (i.n.) aged mice for 6 additional passages (P1-P6) as described above (Fig. 1A). Surprisingly, the RNA copies in the lung reached 10^10.03^ RNA copies/ml at P3, about 250-fold higher than that at P0, and maintained at a similarly high level of RNA copies from P3 to P6 (Fig. 1A).

**Figure 1.**
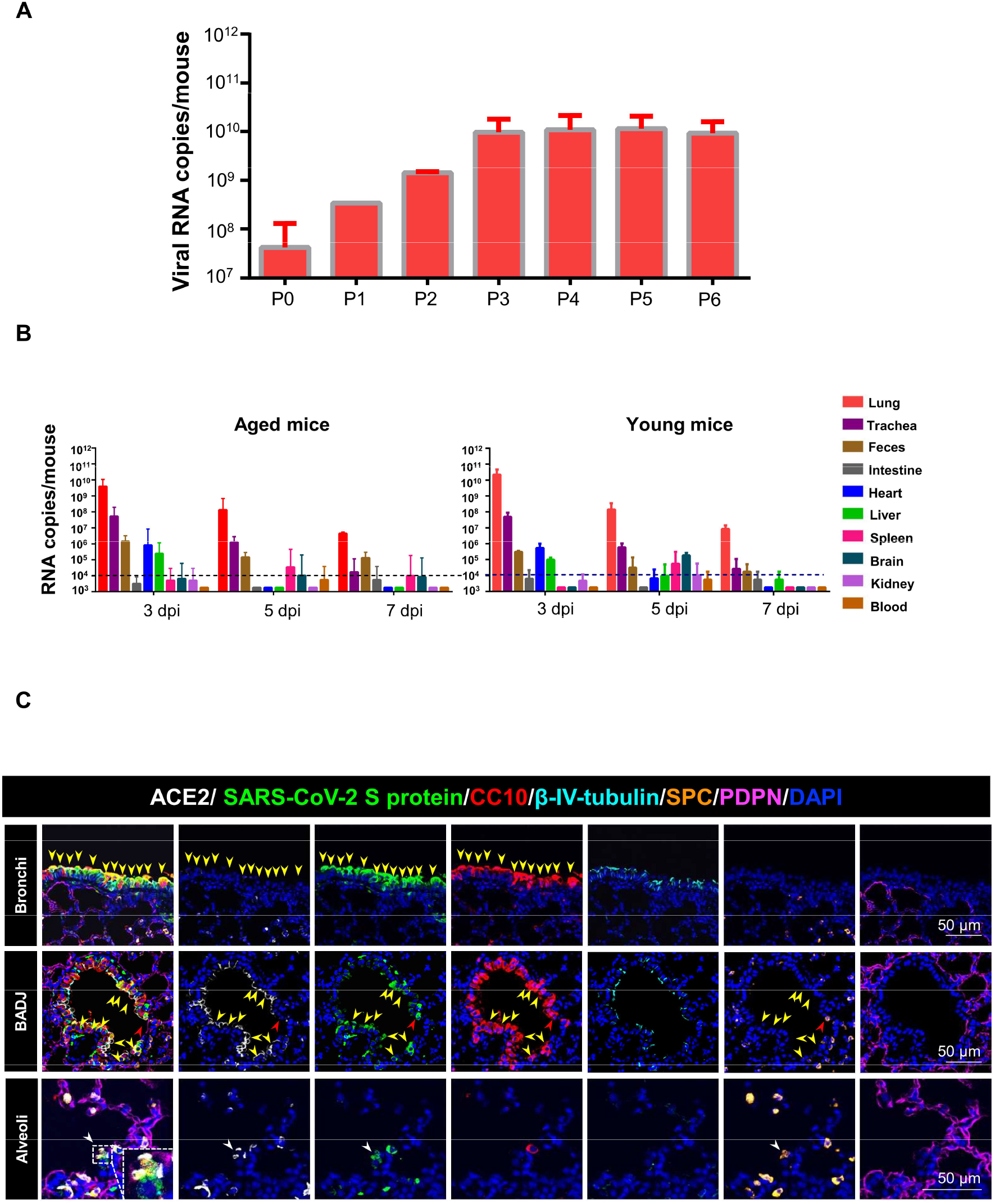
Infection and replication of SARS-CoV-2 at different passage levels in BALB/c mice. (A) Viral RNA copies of mouse-adapted SARS-CoV-2 at passages 0 ~ 6 (P0 ~ P6) were detected by RT-qPCR. Data are presented as means ± standard error of the means (SEM) (n=1-4). (B) Aged (9-month-old) and young (6-week-old) mice were i.n. inoculated with lung homogenates from passage 6 (P6) and sacrificed at 3, 5, and 7 dpi, respectively. Heart, liver, spleen, lung, kidney, brain, intestine, feces, trachea, and sera were collected for detection of viral RNA copies. Dash lines denote the detection limit. Data are presented as means ± SEM (n=3). (C) Immunofluorescence staining for observation of SARS-CoV-2 spike (S) protein (green), ACE2 (white) and cell markers in lung tissues, including CC10 (red), β-IV-tubulin (cyan), PDPN (magenta), and SPC (gold), were shown. Nuclei were shown in blue. The dash box was magnified at the bottom right corner of the figure. Solid yellow arrows indicated SARS-CoV-2^+^/ACE2^+/-^/CC10^+^ cells at the bronchus and bronchioles. Solid red arrows indicated SARS-CoV-2^+^/ACE2^+^/CC10^+^/SPC^+^ cells at the BADJ. Solid white arrows indicated SARS-CoV-2^+^/ACE2^+^/SPC^+^ cells at the alveolus. Scale bars: 50 μm.

We utilized the mouse-adapted SARS-CoV-2 at passage 6 (MASCp6) to infect aged (9-month-old) and young (6-week-old) BALB/c mice. The mice were daily monitored for clinical symptoms and body weight changes, and sacrificed on day 7 post infection. Only the aged mice lost their weight to 5% at 5 days post infection, and then recovered (fig. S1). Results showed a similar level of viral RNA in both aged and young mice at 3, 5, and 7 dpi, respectively, with the peak at 3 dpi (Fig. 1B). The lung had the highest viral load with 10^9.16~10.42^ RNA copies/mouse at 3 dpi of the aged and young mice (fig.S2A). Viral RNA was also detected in trachea, faces, heart, and liver at 3 dpi, and lung, trachea, and faces at 5 and 7 dpi, respectively (Fig. 1B). We then performed an immunofluorescence staining to observe the distribution of viral antigen in the lung tissue. Results revealed robust expression of SARS-CoV-2 S protein along the airways and at the alveolus in both young and aged mice at 3 and 5 dpi, respectively (fig. S2B), whereas no related viral protein was detected in the control mice (data not show). To identify the target cells of this adapted SARS-CoV-2 in mice, we further stained the lung section with different cell markers of lung epithelium, including Club cell (marked by Clara cell 10 kDa protein/CC10), ciliated cell (marked by β-IV-tubulin), alveolar type 1 cell (AT1, marked by podoplanin/PDPN), and alveolar type 2 cell (AT2, marked by surfactant protein C/SPC), in addition to SARS-CoV-2 S protein. As shown in Fig 1C, S protein predominantly colocalized with CC10 and its receptor ACE2 at the bronchus and bronchioles. A few CC10 and SPC double-positive stem cells at the bronchioaveolar-duct junction (BADJ) and SPC-positive AT2 cells in the alveolus were also identified. These findings indicate that Club cells were the major target cells of the mouse-adapted SARS-CoV-2. Collectively, these results suggest that mouse-adapted SARS-CoV-2, MASCp6, can infect and robustly replicate in Club cells and AT2 cells in lung of wild-type BALB/c mice.

### Pathological characteristics of both young and aged BALB/c mice infected with the mouse-adapted SARS-CoV-2

To examine if wild-type young mice are sensitive to the infection of the mouse-adapted SARS-CoV-2, MASCp6, and compare them with the aged mice, we infected young (6-week-old) and aged (9-month-old) BALB/c mice with MASCp6, and at 3, 5 and 7 dpi, respectively, we collected mouse lung to detect tissue damage, and sera to analyze cytokines and chemokines. Histopathological analysis of lung tissues revealed that both aged and young BALB/c mice presented mild to moderate pneumonia after MASCp6 infection (Fig. 2). In the aged mice, lung damage was severe in the infected mice at 3 dpi, with denatured and collapsed epithelial tissues, thickened alveolar septa, alveolar damage, focal exudation and hemorrhage, and activated inflammatory cell infiltration (Fig. 2A). In addition, the endothelial tissues were also denatured, and exhibited adherent inflammatory cells and damage to the basement membrane (Fig. 2A). At 5 dpi, the damage of lung tissue was milder as compared to that on 3 dpi, with denatured epithelial tissues, thickened alveolar septa and lymphatic sheath around vessels (Fig. 2A). In the young mice, the lung damage in the infected mice at 3 dpi was similar to that of the aged mice, with denatured epithelial tissues, thickened alveolar septa, and activated inflammatory cell infiltration (Fig. 2C). At 5 dpi, the epithelial tissues were almost normal, except for the thickened alveolar septa and inflammatory cell infiltration (Fig. 2C). Previous reports on SARS-CoV-2-infected hACE2-Tg mice had mild inflammatory responses and lung damage (*16*). Here we found that MASCp6-infect mice exhibited moderate inflammatory responses with of macrophage infiltration in the lung and increase of some inflammatory cytokines in sera (Fig. 2B and D). Notably, the levels of cytokines, including TNF-α, IL-1β, IL-6, and IL-5, and chemokines, such as MCP-1, G-CSF, GM-CSF, and KC, increased significantly in the aged mice and maintained at a higher level until 5 dpi, compared with the young mice, (fig. S3). To sum up, these results indicate that the mouse-adapted SARS-CoV-2 (MASCp6) efficiently infect both young and old wildtype BALB/c mice, resulting in acute inflammatory responses closely related to the damage of lung tissues.

**Figure 2.**
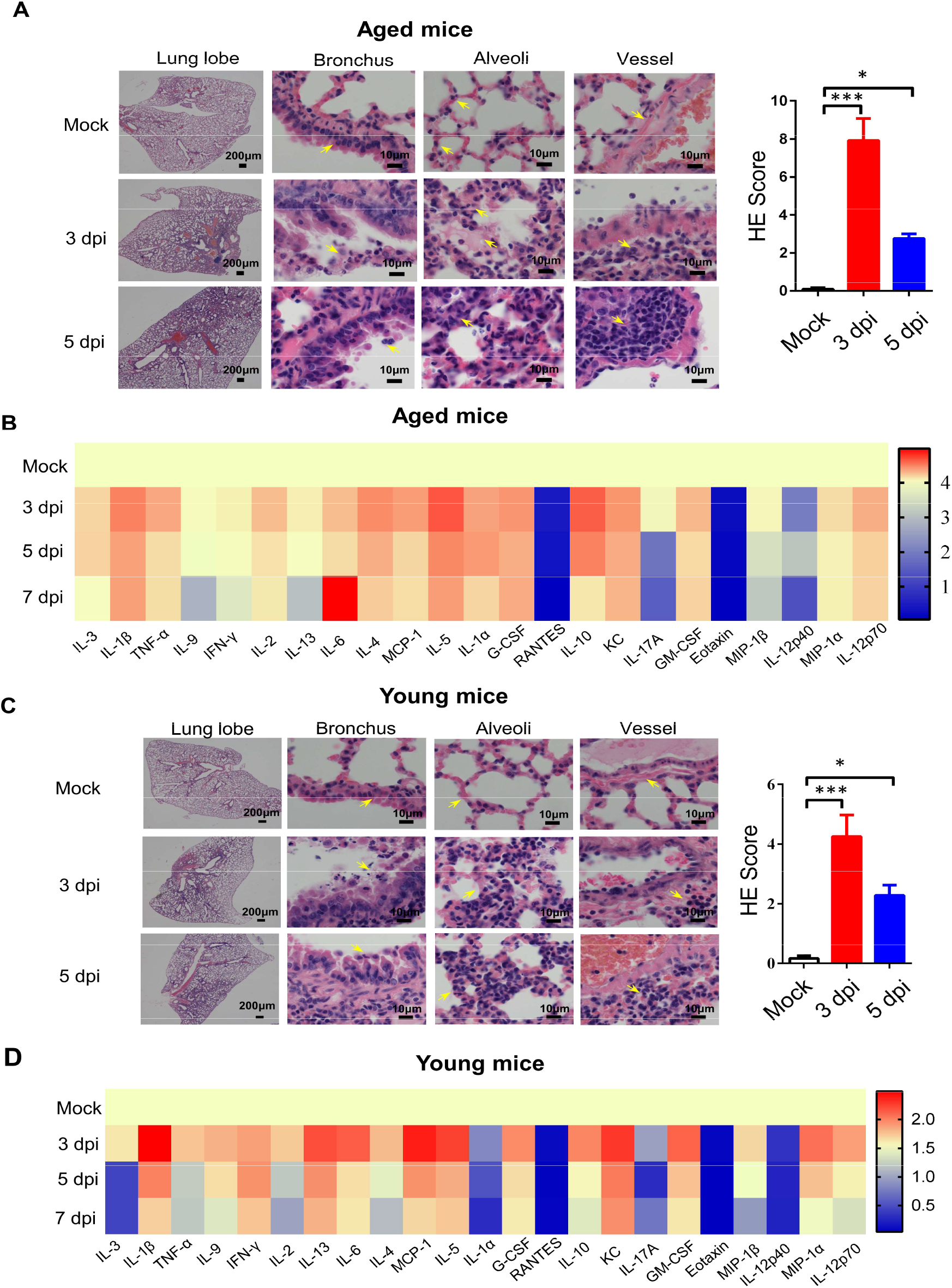
Analysis of histopathological changes and inflammatory response in young and aged BALB/c mice infected with adapted SARS-CoV-2. Aged (9-month-old) and young (6-week-old) naïve BALB/c mice were i.n. infected with 30 μl of mouse-adapted SARS-CoV-2, MASCp6. Lung tissues were removed for analysis of histopathological damage, and sera were collected for detection of cytokines at the indicated time points. (A, C) Hematoxylin and eosin (H&E) staining of tissue sections of lung lobe, bronchus, alveoli, and vessel from mice infected with SARS-CoV-2 MASCp6 at 3 and 5 dpi (yellow arrow shows damage). Semi-quantitative analysis of histopathological changes of lung tissues of infected mice at 3 and 5 dpi. Data are presented as means ± SEM (n=3). (B, D) Cytokine and chemokine production by Luminex assay in sera of mice infected with SARS-CoV-2 MASCp6 at 3, 5, and 7 dpi, respectively. Data are presented as means ± SEM (n=5). Statistical significance was analyzed by unpaired Student’s t tests. n=5. \**P* < 0.01, \*\**P* < 0.01.

### Identification of a key mutation in mouse-adapted SARS-CoV-2 with infectivity

To investigate potential mutations in MASCp6 associated with increased infectivity, we compared the genomic sequences between MASCp6 and the original wild-type SARS-CoV-2 strain (BetaCoV/Wuhan/AMMS01/2020). Among a total of five mutations identified in MASCp6, four of them resulted in the change of amino acid residues (Fig. 3A). These mutations were localized at the open reading frame 1ab (ORF1ab), S, and nucleocapsid (N) genes. The mutation in the S gene occurred at the RBD region, leading to the change of residue 501 of S protein from asparagine to tyrosine (N501Y). SARS-CoV-2 binds its receptor ACE2 through the RBD, and residue N501 at the RBD has been identified as one of the five key residues responsible for the receptor recognition and host range of SARS-CoV-2(*18*). Protein structure and affinity prediction analysis indicated that N501Y mutation could increase the binding affinity between mouse ACE2 (mACE2) and RBD of SARS-CoV-2 S protein (Fig.3B). The binding affinity between MASCp6 and mACE2 was also confirmed by immunofluorescence straining, which showed strong signals of viral S protein and mACE2 (Fig. 3C). In sum, these data indicate that the increased infectivity by adapted SARS-CoV-2 (MASCp6) in mice might be due to the enhanced binding affinity between its mutant RBD containing N501Y mutation and mACE2 receptor.

**Figure 3.**
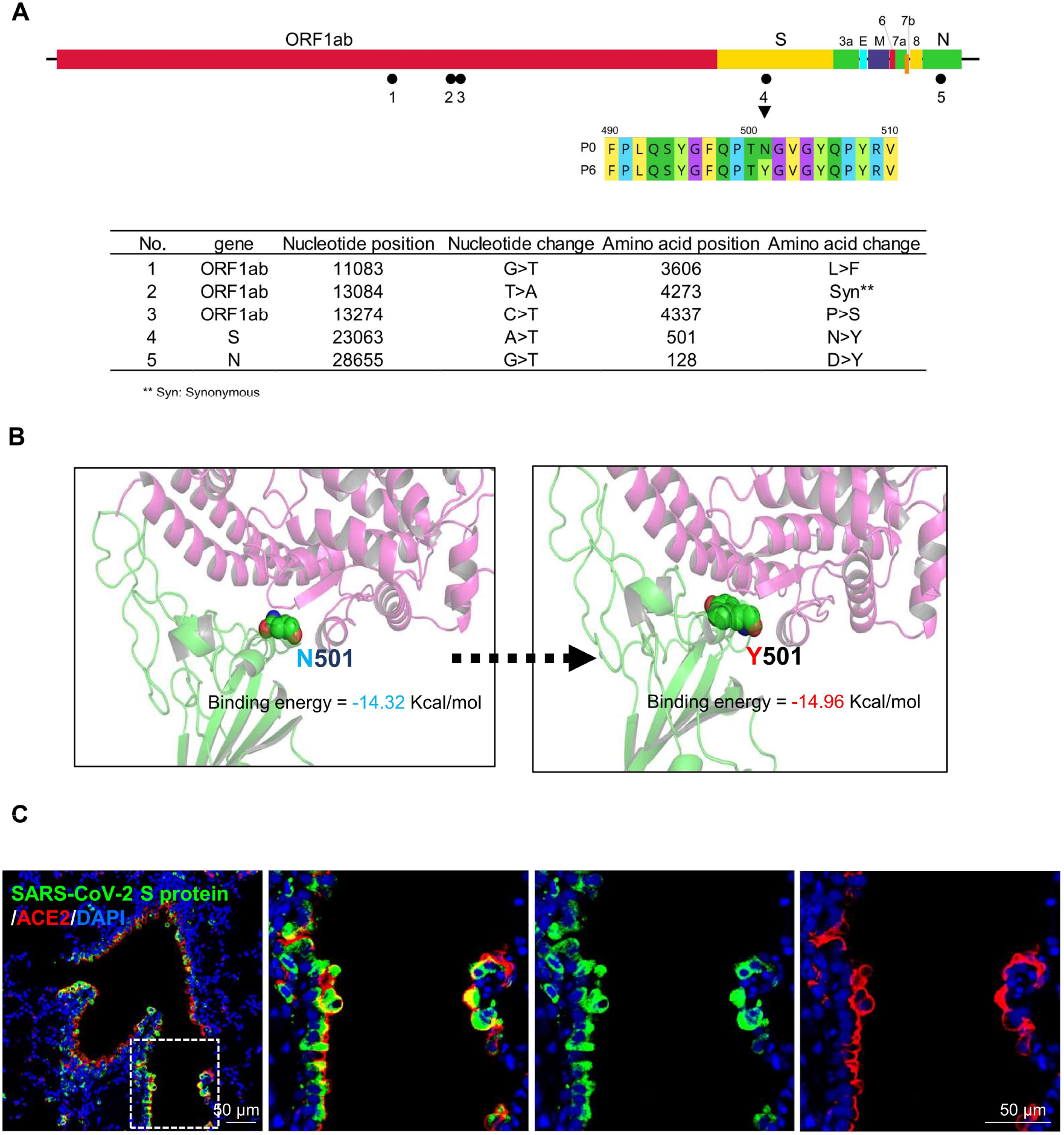
Mutations identified in the mouse-adapted SARS-CoV-2 strain MACSp6. (A) Schematic diagram of the SARS-CoV-2 genome, indicating the mutations identified in the mouse-adapted SARS-CoV-2. (B) RBD in the Spike structure is shown as a green helix. Homology modeling of mACE2, in which the helix is colored pink. The binding energy between S–ACE2 interaction is improved from −14.32 Kcal/ mol (N501) to −14.96 Kcal/mol (Y501). (C) Colocalization of SARS-CoV-2 S protein (green) and mouse ACE2 (red) in lung tissues (n=3). The dash box was magnified in the panels at the right side. Scale bars: 50 μm.

### The protective efficacy of RBD-based SARS-CoV-2 subunit vaccine in the established mouse model

To identify potential application of this mouse-adapted SARS-CoV-2 infection model for vaccine evaluation, we constructed a subunit vaccine, RBD-Fc, consisting of SARS-CoV-2 S protein RBD fused with a human IgG Fc (fig. S4, A and B) (*19*). RBD-Fc was highly expressed in an established CHO stable cell line and purified with high purity (fig. S4B, right). It bound strongly to hACE2 in hACE2/293T cells (fig. S4C), indicating its good functionality.

To investigate the immunogenicity of this vaccine, we intramuscularly (i.m.) immunized BALB/c mice (6-8 week-old) with RBD-Fc protein (10 μg/mouse) or PBS (control) in the presence of aluminum hydroxide (100 μg/mouse) adjuvant, and boosted once at 2 weeks, followed by collection of sera for detection of antibodies, including neutralizing antibodies. As shown in Fig. 4A, RBD-Fc elicited SARS-CoV-2 RBD-specific IgG antibody response 2 weeks after 1^st^ immunization, and the antibody titer increased rapidly at 2 weeks after 2^nd^ immunization, which maintained at the similarly high level for up to 4 weeks. Importantly, these antibodies potently neutralized live SARS-CoV-2 infection (Fig. 4B).

**Figure 4.**
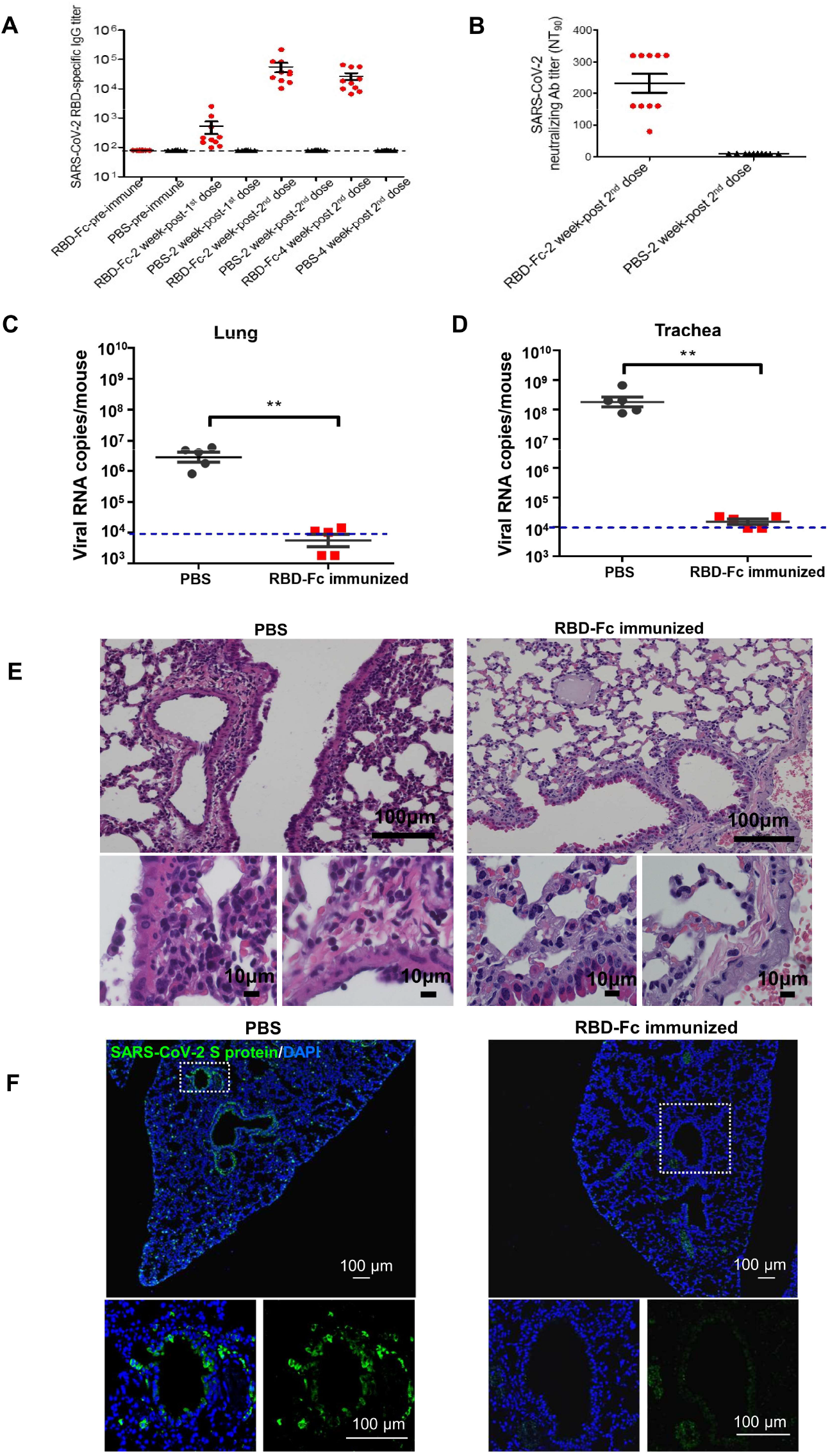
SARS-CoV-2 RBD subunit vaccine elicited high titers of neutralizing antibodies that protected immunized mice against SARS-CoV-2 challenge. BALB/c mice were intramuscularly (i.m.) vaccinated with RBD-Fc or PBS control plus aluminum adjuvant and boosted once at 2 weeks. Sera were collected before immunization, 2 weeks after the 1^st^ immunization, and 2 and 4 weeks, respectively, after the 2^nd^ immunization for detection of SARS-CoV-2 RBD-specific IgG antibody and/or neutralizing antibodies. Four weeks after the 2^nd^ immunization, mice were i.n. inoculated with the mouse-adapted SARS-CoV-2 MACSp6, and their tissues were collected for detection of viral load and lung pathology at 5 dpi. (A) SARS-CoV-2 RBD-specific IgG antibody detected at the indicated time points (n=10). (B) SARS-CoV-2 neutralizing antibody titer (n=10). (C-D) Viral RNA detection in lung and trachea of both immunized mice and control mice infected with the mouse-adapted SARS-CoV-2 (n=5). (E) H&E staining. Focal perivascular and peribronchiolar inflammation and thickened alveolar septa were observed in the lung of mice receiving PBS, but no obvious lung damage in the RBD-Fc-immunized mice was noticed. (F) Immunofluorescence staining of mouse lung paraffin sections for adapted-SARS-CoV-2 S protein (green) and DAPI (blue) (n=3). The dotted boxs were magnified in the panels at the bottom of the figures. Scale bars: 100 μm.

To detect the protective efficacy of this vaccine in our established mouse model, we challenged mice with the adapted SARS-CoV-2, MASCp6 (Infectious virus titer of 10^8.81^ RNA copies /mouse) four weeks after 2nd immunization, and detected for viral replication and histopathological changes 5 days after challenge. Results indicated that RBD-Fc-immunized mice had undetectable viral RNA copies in their lung and trachea, the major organs for SARS-CoV-2 replication, whereas control mice had high viral RNA copies in these tissues (Fig. 4C and D). These results were confirmed by immunofluorescence staining, showing strong signal of SARS-CoV-2 S protein in the lung tissue of control mice, but not in the related tissue of the vaccinated mice (Fig. 4F). In addition, there was no lung damage in the RBD-Fc-immunized mice, whereas inflammatory lung injury with focal perivascular and peribronchiolar inflammation and thickened alveolar septa was observed in the lung tissue of control mice (Fig. 4E). In conclusion, the above data suggest that immunization of RBD-Fc subunit vaccine is able to fully protect the immunized mice against SARS-CoV-2 challenge and that the established mouse model can be effectively used to evaluate the protective efficacy of SARS-CoV-2 vaccines.

## Discussion

SARS-CoV-2 is a novel CoV that has not been previously identified in humans. SARS-CoV-2-caused COVID-19 is spreading globally with no available countermeasures to stop this severe pandemic. Moreover, no convenient and economic small animal models are currently available to evaluate the protective efficacy of anti-SARS-CoV-2 vaccines and therapeutics.

An ideal animal model for SARS-CoV-2 infection should be susceptible to viral infection and replication in respiratory tissues with histopathological presentations of COVID-19. An economical, nontransgenic inbred small animal model could provide a large number of animals for repeated experiments to generate reproducible data for statistical analysis. Because of their easy handling and breeding, mice have been considered as the mainstay for biomedical research. We know that mouse DPP4 does not support MERS-CoV infection (*20*), and that mouse ACE2 does not support SARS-CoV-2 infection. Mice transgenic for hACE2 have been used to develop a SARS-CoV-2 infection animal model, and it presents mild disease (not lethal) with an RNA peak of 10^6.77^ copies/ml or 10^2.44^ TCID_50_/100 μl at 3 dpi (*16*). Ferrets and cats are permissive to SARS-CoV-2 infection with the highest infectious virus titer of 10^1.53^ RNA copies/g or 10^1.4^ TCID_50_/g in the lung of infected ferrets at 4 dpi (*21, 22*). Golden or Syrian hamsters are also susceptible to SARS-CoV-2 infection with viral load as high as 10^5^-10^7^ TCID_50_/g in the lung and progressive exudative inflammation with weight loss, lethargy, and rapid breathing at 2 dpi, while they can recover from infection about one week post-infection (*23*). Except for the above models, nonhuman primates (NHPs), which are closest to humans phylogenetically, have been used to develop a SARS-CoV-2 infection model. SARS-CoV-2-infected macaques showed no, or mild, clinical symptoms with viral antigens detected mostly in the nasolacrimal system (*24, 25*). Notably, the short supply and high cost of hACE2-transgenic mice, as well as the costs involved in purchasing and handling other animals permissive to SARS-CoV-2, including NHPs, make it very difficult for researchers to evaluate the *in vivo* protective efficacy of anti-SARS-vaccines, antibodies and therapeutics.

Different from the above models, here we have developed a convenient, economic, available, and effective mouse model for SARS-CoV-2 infection. SARS-CoV-2 was rapidly adapted in the inbreed aged (9-month-old) BALB/c mice, resulting in generation of very high viral load (10 ^9.16-10.42^ RNA copies/mouse) because of stable viral replication in the lungs. The SARS-CoV-2 MASCp6 strain adapted in the aged mice could also efficiently infect young (6-to 8-month-old) BALB/c mice and C57BL/6 mice (data not shown) with high viral RNA copies in lungs and tracheas. Our data suggest that this mouse-adapted SARS-CoV-2 strain can infect at least two major mouse strains (BALB/c and C57BL/6) at various ages. Deep sequencing of genes of this mouse-adapted SARS-CoV-2 showed five mutations absent in wild-type human SARS-CoV-2, one of which occurred at residue 501 (N501Y) of RBD in the S protein of SARS-CoV-2. This mutation was predicted to increase binding affinity to mACE2, potentially contributing to broad infectivity of the adapted MASCp6 strain in mice.

Histopathological findings from the adapted SARS-CoV-2-infected BALB/c mouse model resembled the manifestations of mild to moderate acute clinical cases. SARS-CoV-2 infection was seen in the airway of this mouse model, presenting denatured trachea, changes of inflammation in pulmonary alveoli with detection of viral antigen in trachea, bronchiole and some type II pneumocytes. The accelerated systemic and local immune responses, such as elevated inflammatory cytokines and chemokines, in sera and lung macrophages of infected mice were similar to those observed in COVID-19 patients(*26*). Although the mice were not lethal after 6 passages of SARS-CoV-2 infection, they had stable viral replication and high viral load in lungs, and moderate lung damage.

Using this well-developed mouse-adapted SARS-CoV-2 infection animal model, here we evaluated the in vivo protective efficacy of an RBD-based SARS-CoV-2 subunit vaccine. We have previously demonstrated that RBD in the S protein of Coronaviruses, including SARS-CoV and MERS-CoV, is the key target for development of vaccines and therapeutics (*13, 14, 19, 27, 28*). Here we showed that the RBD-based SARS-CoV-2 vaccine elicited potent antibody responses with strong neutralizing activity against SARS-CoV-2 infection and full protection of the vaccinated mice against infection of the mouse-adapted SARS-CoV-2 strain (MASCp6) with no detectable viral RNA in the lung and trachea, two major organs with high susceptibility to SARS-CoV-2 infection (*29*).

Overall, our established mouse-adapted SARS-CoV-2 infection mouse model can be conveniently, economically, and effectively used for evaluation of the in vivo protective efficacy of SARS-CoV-2 vaccines as well as anti-SARS-CoV-2 antibodies and therapeutics. Moreover, the RBD-based SARS-CoV-2 subunit vaccine tested here has great promise for further development as an effective vaccine to prevent SARS-CoV-2 infection and contain COVID-19 pandemics.

## Acknowledgements

We thank Dr. Xue-Dong Yu for excellent technical and biosafety support. H.G.,Q.C.,G.Y.,L.H.,H.F.,Y.D.,Y.W.,Y.T.,Z.Z.,Y.C.,Y.L.,X.L.,J.L.,N.Z.,X.Y.,S.C.,G.Z.,G.H.,D. L. performed experiments; X.W.,H.W.,X.Y.,Y.L.,Y.H.,Z.X.,S.G., and X.S. analyzed data; S.J., S.S.,C.Q., and Y.Z. designed the study and wrote the manuscript. This work was supported by the National Natural Science Foundation of China (No.82041006 and 82041025), The National Program of Infectious Diseases Fund (No.2017ZX10304402-003), the National Science and Technology Major Project of China (No.2018ZX09711003). All the authors declare no competing interests. C.F.Q. was supported by the National Science Fund for Distinguished Young Scholar (No.81925025), and the Innovative Research Group (No.81621005) from the NSFC, and the Innovation Fund for Medical Sciences (No.2019-I2M-5-049) from the Chinese Academy of Medical Sciences.

## Materials and Methods

### Ethics statement

All procedures involving cells and animals were conducted in Biosafety Level 3 laboratory (BSL-3) and approved by the Animal Experiment Committee of Laboratory Animal Center, Beijing Institute of Microbiology and Epidemiology (approval number: IACUC-DWZX-2020-002). Animal studies were carried out in strict accordance with the recommendations in the Guide for the Care and Use of Laboratory Animals.

### Virus and cell

SARS-CoV-2 strain BetaCoV/Wuhan/AMMS01/2020 was originally isolated with Vero cells from a patient returning from Wuhan. The virus was amplified and titrated by standard plaque forming assay on Vero cells. All experiments involving infectious SARS-CoV-2 were performed in biosafety level 3 (BSL-3) containment laboratory in Beijing Institute of Microbiology and Epidemiology.

### Serial passage of SARS-CoV-2 in aged BALB/c mice

Adaptation of SARS-CoV-2 was achieved by serial passage through lungs of BALB/c mice. A dose of 7.2×10^5^ plaque forming unit (PFU) of SARS-CoV-2 (BetaCoV/Wuhan/AMMS01/2020) was administered intranasally (i.n.) to three anesthetized, 9-month-old female BALB/c mice in a total volume of 40 μL. Three days after inoculation, the mice were euthanized, and the lung of each mouse was removed and homogenized as a 10% w/v suspension in Dulbecco’s Modified Eagle’s Medium (Invitrogen) containing 100 IU/ml Penicillin (Gibco), 0.1 mg/ml Streptomycin (Gibco), and 10% Fetal Bovine Serum (FBS). The lung homogenate was clarified by centrifugation at 6,000 rpm for 6 min, and the supernatant of lung homogenates from three mice was administered i.n. into three naïve mice after filtration and mixing completely. The process of i.n. inoculation of three female aged BALB/c mice was repeated 6 times. After detection of SARS-CoV-2 RNA in the lung homogenate, and viral load was detected by cytopathic effect (CPE) and plaque assay, respectively. Virus from passage 6, named MASCp6, was stored and aliquot for the following studies.

### Measurement of viral RNA

Tissue homogenates were clarified by centrifugation at 6,000 rpm for 6 min, and the supernatants were transferred to a new EP tube. Viral RNA (vRNA) was extracted using the QIAamp Viral RNA Mini Kit (Giagen) according to the manufacturer’s protocol. vRNA quantification in each sample was performed by quantitative reverse transcription PCR (RT-qPCR) targeting the S gene of SARS-CoV-2. RT-qPCR was performed using One Step PrimeScript RT-PCR Kit (Takara, Japan) with the following primers and probes: CoV-F3 (5’-TCCTGGTGATTCTTCTTCAGGT-3’); CoV-R3 (5’-TCTGAGAGAGGGTCAAGTGC-3’); and CoV-P3 (5 ‘-AGCTGCAGCACCAGCTGTCCA-3 ‘).

### RNA extraction and genome sequencing

Total RNA from the lung homogenate was extracted using High Pure Viral RNA Kit (Roche, Switzerland), and the purified vRNA was used to synthesize the first-strand cDNA by reverse transcription using SuperScript VILO cDNA Synthesis Kit (ThermoFisher, USA) according to the manufacturer’s protocol. The sequencing library was constructed using Ion Ampliseq Library Kit 2.0 (ThermoFisher), and sequenced on an Ion Torrent S5Plus sequencer (ThermoFisher). A total of 230,225 reads were produced. The average length of reads was 221 nucleotide (nt).

### Genome assembly and variation identification

Sequences were assembled and analyzed with CLC Genomic Workbench (Qiagen, Germany). All reads were mapped to SARS-CoV-2 reference genome (Wuhan-Hu-1, GenBank accession number MN908947). The consensus sequence was extracted. When a variation was identified after comparison of the genome sequences, independent RT-PCR reactions were run, and subsequent RT-PCR products were sequenced through the region containing the putative mutation to confirm the mutant residues.

### Genome deposition

The genome sequence of mouse-adapted SARS-CoV-2 had been uploaded to the Genome Warehouse in National Genomics Data Center, Beijing Institute of Genomics (BIG), Chinese Academy of Sciences, with the accession number of GWHACFH00000000 that is publicly accessible at https://bigd.big.ac.cn/gwh.

### Histopathological analysis

Female BALB/c mice at 6 weeks old and 9 months old were maintained in a pathogen-free facility and housed in cages containing sterilized feed and drinking water. Following intraperitoneal (i.p.) anesthetization with sodium pentobarbital, all mice were i.n. inoculated with 30 μL of the adapted SARS-CoV-2, MASCp6. Lung and trachea tissues were removed, and paraffin-embedded in accordance with the standard procedure. Sections at 5 μm thickness were stained with hematoxylin and eosin (H&E), and examined by light microscopy. Lung tissue lesions were assessed according to the extent of denatured and collapsed bronchiole epithelial cells, degeneration of alveoli pneumocytes, infiltration of inflammatory cells, edema, hemorrhage, exudation and expansion of parenchymal wall. The semiquantitative assessment was scored for the severity of lung damage according to the above criteria (0, normal; 1, mild; 2, moderate; 3, marked). The cumulative scores of the severity provided the total score per animal, and the average of three to four animals from each group was taken as the total score for that group.

### Multiplex immunofluorescent assay

5 μm paraffin sections were deparaffinized in xylene and rehydrated in a series of graded alcohols. Antigen retrievals were performed in citrate buffer (pH6) with a microwave (Sharp, R-331ZX) for 20-min at 95°C followed by a 20-min cool down at room temperature. Multiplex fluorescence labeling was performed using TSA-dendron-fluorophores (NEON 7-color Allround Discovery Kit for FFPE, Histova Biotechnology, NEFP750). Briefly, endogenous peroxidase was quenched in 3% H_2_O_2_ for 20-min, followed by blocking reagent for 30-min at room temperature. Primary antibody was incubated for 2-hr in a humidified chamber at 37°C, followed by detection using the HRP-conjugated secondary antibody and TSA-dendron-fluorophores. Afterwards, the primary and secondary antibodies were thoroughly eliminated by heating the slides in retrieval/elution buffer (Abcracker^®^, Histova Biotechnology, ABCFR5L) for 10-sec at 95°C using microwave. In a serial fashion, each antigen was labeled by distinct fluorophores. Multiplex antibody panels applied in this study are: ACE2 (Abcam, ab108252, 1:200), SARS-CoV Spike (Sinobiological, 40150-T52, 1:2000); β-IV-tubulin IV (Abcam, ab179504, 1:1000), CC10 (Millipore, 07-623, 1:500), Podoplanin (Sinobiological, 50256-R066, 1: 1000), SP-C (Abcam, ab211326, 1:500). After all the antibodies were detected sequentially, the slices were imaged using the confocal laser scanning microscopy platform Zeiss LSM880.

### Cytokine and chemokines analysis

Cytokines and chemokines in mouse sera were measured using Bio-Plex Pro Mouse Cytokine Grp I Panel 23-Plex (BIO-RAD). A panel of inflammatory cytokines and chemokines was detected according to the manufacturer’s protocol. The data were collected on Luminex 200, and analyzed by Luminex PONENT (Thermo Fisher).

### Construction, expression and purification of recombinant protein

The code optimized genes encoding the fragment containing 194 aa (331–524) of the SARS-CoV-2 S protein RBD region with human IgG1 Fc was inserted into the GS Gene Expression Vector. The constructed recombinant plasmid (RBD007) was confirmed by sequencing analysis. Briefly, the recombinant RBD007 plasmid was transfected using FuGENE 6 transfection reagents (Roche Applied Science, Indianapolis, IN) into CHO-K1 cells precultured in F-12K medium (American Type Culture Collection, Manassas, VA), according to the manufacturer’s instructions. Stable expressing cells were selected by culturing in selective medium. The expression and purification of the RBD-hFc protein were performed in GLP laboratory. The recombinant RBD-hFc protein was purified through affinity chromatography and anion exchange chromatography. The purity of recombinant protein were detected by SDS-PAGE and HPLC.

### Flow cytometry analysis

Flow cytometry analysis was carried out to detect the binding of SARS-CoV-2 RBD-Fc to hACE2 receptor in hACE2-expressing 293T (hACE2/293T) cells. Briefly, hACE2/293T cells were incubated with SARS-CoV-2 RBD-Fc protein or Fc control (10 μg/ml) for 20 min at 37°C. After staining with FITC-conjugated goat anti-human IgG antibody (1:500; Thermo Fisher Scientific), the cells were analyzed by flow cytometer (BD LSRFortessa™) and Flowjo software.

### Animal immunization and challenge studies

BALB/c mice were immunized with SARS-CoV-2 RBD-Fc protein. Briefly, mice (6-to 8-week-old) were intramuscularly (i.m.) vaccinated with RBD-Fc (10 μg /mouse) or PBS control in the presence of aluminum (100 μg/mouse) adjuvant and boosted with the same immunogen and adjuvant at 2 weeks. Sera were collected before immunization and 2 or 4 weeks after each immunization to detect SARS-CoV-2 S-specific antibody responses and/or neutralizing antibodies as described below. Four weeks after the boosted immunization, mice were inoculated intranasally with the adapted SARS-CoV-2, MASCp6 (infectious virus titer of 10^8.81^ RNA copies /mouse). All mice were observed daily. On day 5 postinfection, seven mice in each group were sacrificed, and their lung tissues were removed for detection of viral load. Lungs in the other three mice were embedded for pathological analysis as described above.

### ELISA

ELISA was performed to detect SARS-CoV-2 RBD-specific IgG antibodies in the immunized mouse sera. Briefly, ELISA plates were precoated with SARS-CoV S1 (1 μg/ml) overnight at 4°C and blocked with 2% fat-free milk in PBST for 2 h at 37°C. Serially diluted sera were added to the plates and incubated for 2 h at 37°C. After four washes, the bound antibodies were detected by incubation with horseradish peroxidase (HRP)-conjugated anti-mouse IgG antibody (1:5,000, Thermo Fisher Scientific) for 1 h at 37°C. The reaction was visualized by addition of substrate 3,3’,5,5’-Tetramethylbenzidine (TMB) (Sigma, St. Louis, MO) and stopped by H_2_SO_4_ (1N). The absorbance at 450 nm was measured by an ELISA plate reader (Tecan, San Jose, CA).

### SARS-CoV-2 neutralization assay

A micro-neutralization assay was carried out to detect neutralizing antibodies against SARS-CoV-2 infection. Briefly, mouse sera at 2-fold serial dilutions were incubated with SARS-CoV-2 (100 TCID_50_) for 1 h at 37°C and added to Vero cells. The cells were observed daily for the presence or absence of virus-induced CPE and recorded at 3 dpi. Neutralizing antibody titers were determined as the highest dilution of sera that completely inhibited virus-induced CPE in at least 90% of the wells (NT_90_).

### Statistical analysis

Statistical analyses were carried out using Prism software (GraphPad). All data are presented as means ± standard error of the means (SEM). Statistical significance among different groups was calculated using the Student’s *t* test, Fisher’s Exact test, or Mann-Whitney test *, **, and *** indicate *P* < 0.05, *P* < 0.01, and *P* < 0.001, respectively.

---

**Supplemental Figure 1.**
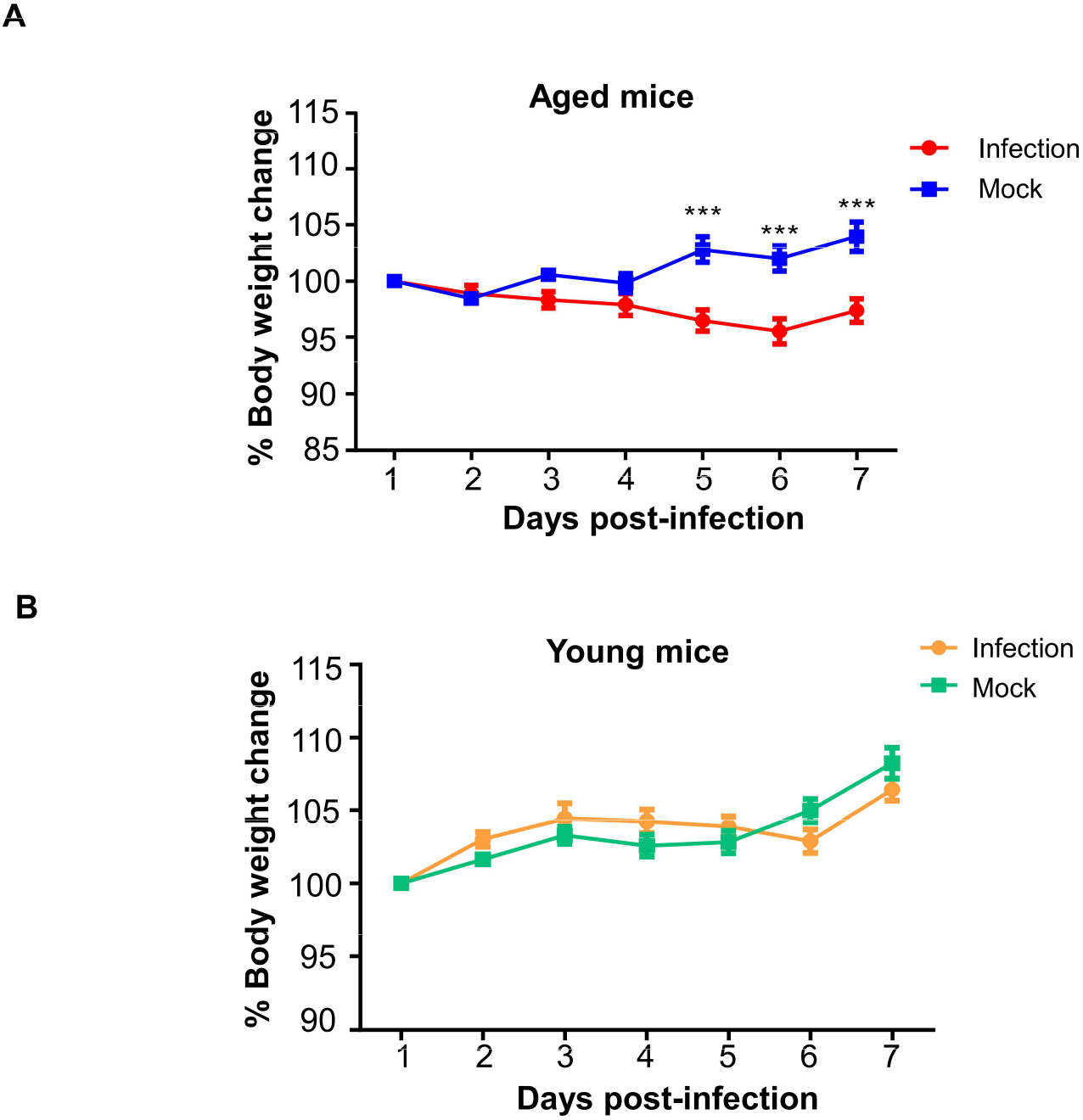
Body weight change of mice infected with MASCp6. Aged (9-month-old) and young (6-week-old) naïve BALB/c mice (n=6) were i.n. infected with MASCp6 and their body weight change was monitored daily.

**Supplemental Figure 2.**
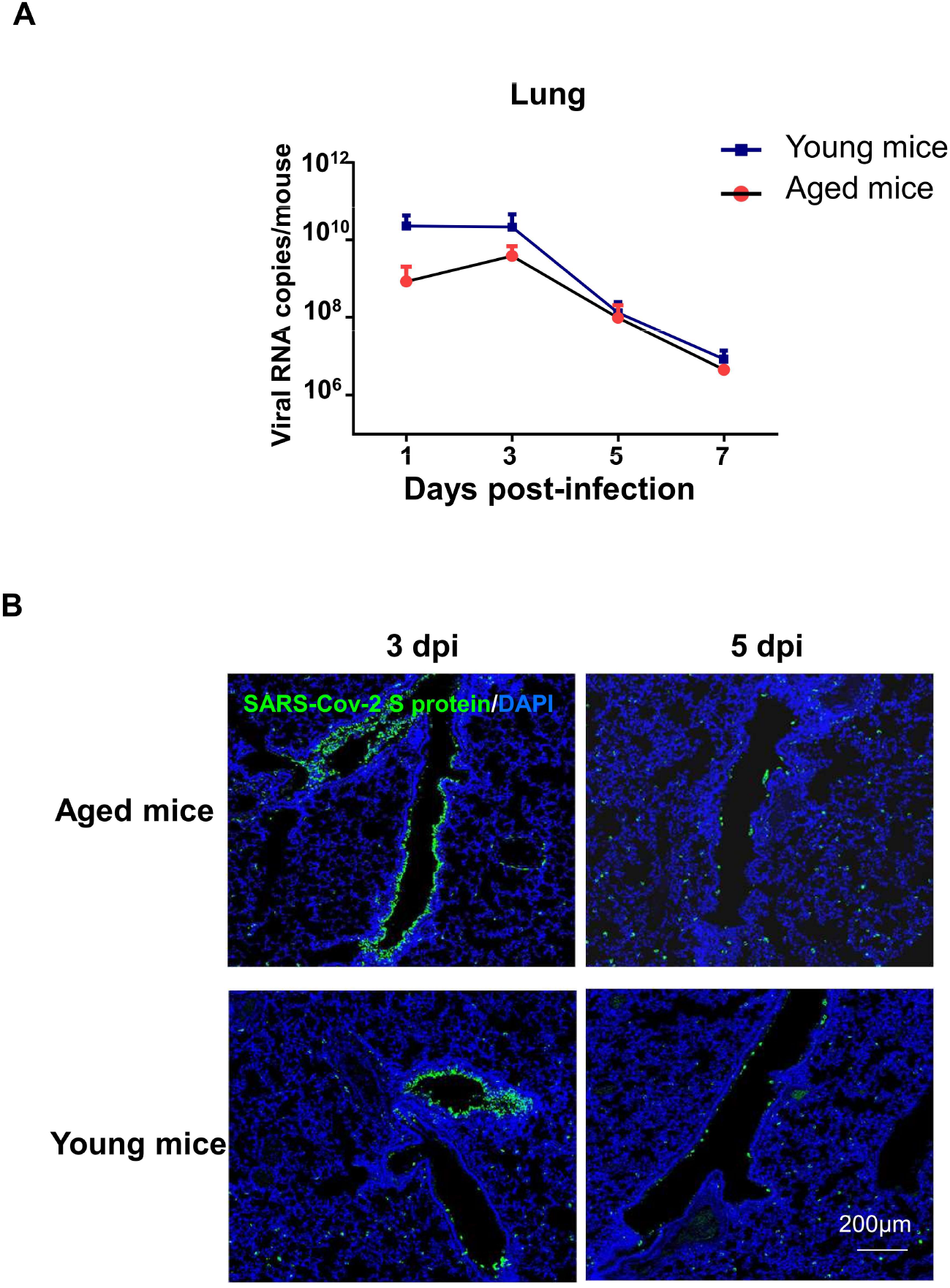
Viral replication in BABL/c infected with MASCp6. (A) Viral RNA copies in lung of mice infected with MASCp6. Aged (9-month-old) and young (6-week-old) naïve BABL/c mice (n=3) were i.n. infected with lung homogenates collected at passage 6 and detected viral RNA copies in lung was detected at 1, 3, 5, 7 pdi, respectively, via RT-qPCR. (B) Immunofluorescence staining for the viral location in lung tissue. Scale bars: 200 μm.

**Supplemental Figure 3.**
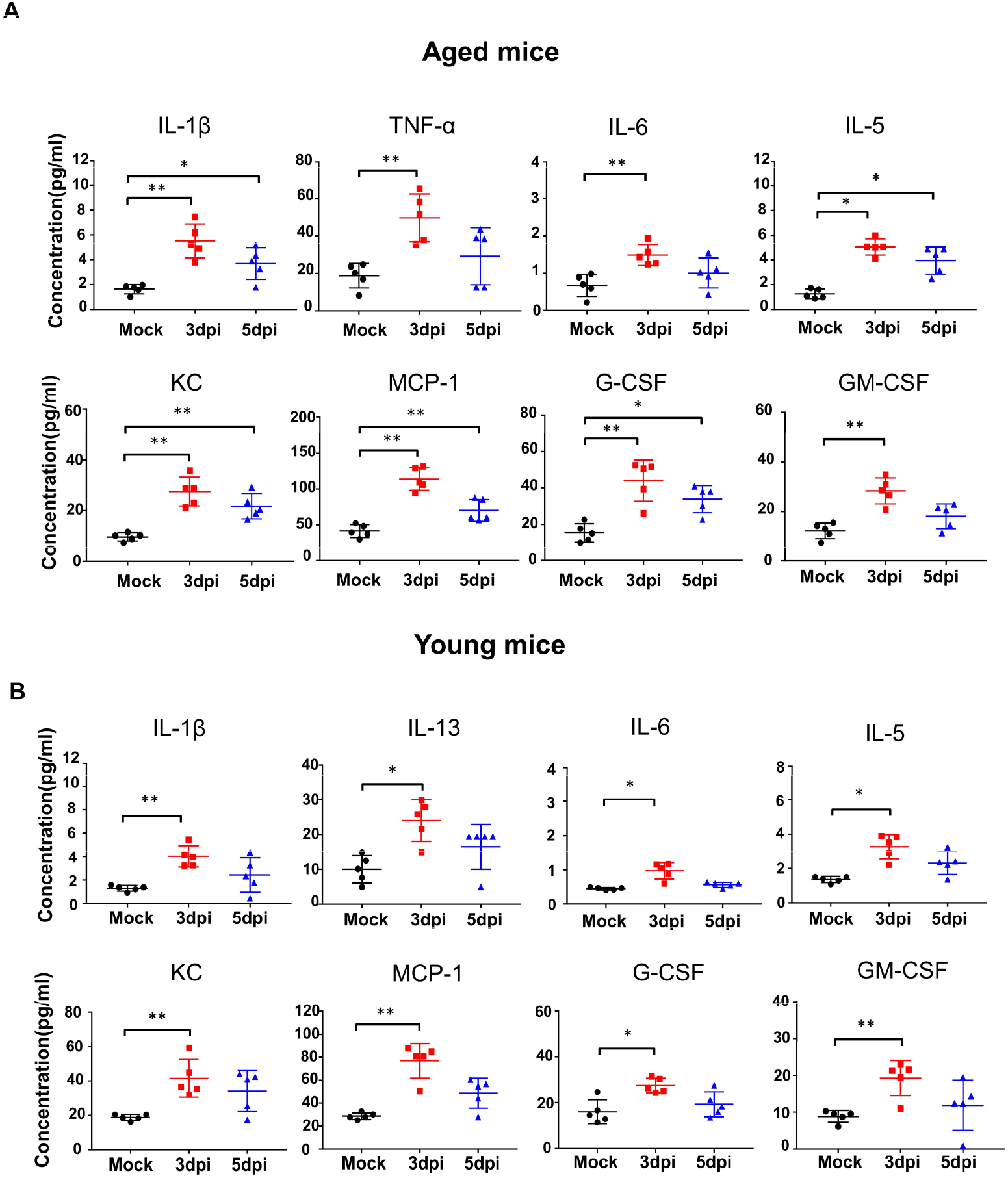
Cytokines and chemokines production in the sera of MASCp6-infected mice. The level of cytokines and chemokines in the sera of aged (A) and young (B) mice at 3 and 5 dpi, respectively. Statistical significance was analyzed by unpaired Student’s t tests. n=5. *P < 0.01, ** *P* < 0.01.

**Supplemental Figure 4.**
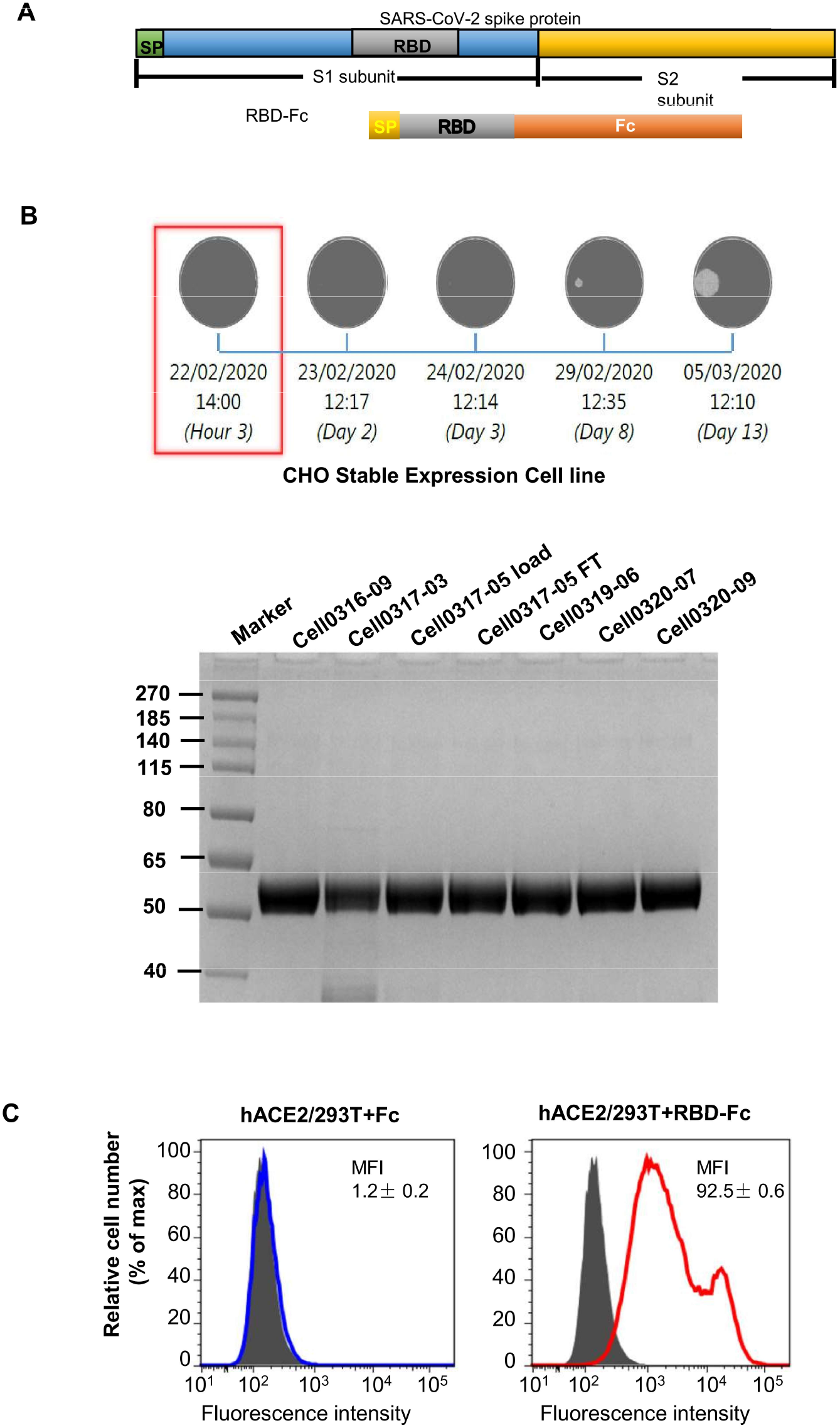
Construction and characterization of RBD-based SARS-CoV-2 subunit vaccine. (A).Schematic diagram of design of RBD-based SARS-CoV-2 subunit vaccine. (B) Selection of stable expressing cell lines and purification of the recombinant RBD-hFc protein of SARS-CoV-2. (C) The bounding ability of the recombinant RBD-hFc protein to hACE2 in hACE2/293T cells.

